# Cardiomyocyte antisense transcription regulates exon usage in the elastic spring region of Titin to modulate sarcomere function

**DOI:** 10.1101/2024.07.10.602998

**Authors:** Selvi Celik, Ludvig Hyrefelt, Tomasz Czuba, Yuan Li, Juliana Assis, Oscar André, Joakim Sandstedt, Pontus Nordenfelt, Kristina Vukusic, J. Gustav Smith, Olof Gidlöf

**Author notes:** Corresponding author contact information: Dr. Olof Gidlöf, Lund University, Dept. of Cardiology, BMC D12, Solvegatan 19, SE-221 84 Lund.

## Abstract

**Background:** The spring-like sarcomere protein Titin (TTN) is a key determinant of cardiac passive stiffness and diastolic function. Alternative splicing of TTN I-band exons produce protein isoforms with variable size and elasticity, but the mechanisms regulating TTN exon skipping and isoform composition in the human heart are not well studied. Non-coding RNA transcripts from the antisense strand of protein-coding genes have been shown to regulate alternative splicing of the sense gene. The *TTN* gene locus harbours >80 antisense transcripts with unknown function in the human heart. The aim of this study was to determine if *TTN* antisense transcripts play a role in alternative splicing of *TTN*.

**Methods:** RNA-sequencing and RNA in situ hybridization (ISH) of cardiac tissue from unused organ donor hearts (n=7) and human induced pluripotent stem cell-derived cardiomyocytes (iPS-CMs) were used to determine the expression and localization of TTN antisense transcripts. The effect of siRNA-mediated knock down of *TTN* antisense transcripts on *TTN* exon usage in iPS-CMs was determined using RNA-sequencing. Live cell imaging with sarcomere tracking was used to analyze the effect of antisense transcript knock down on sarcomere length, organization and contraction dynamics. RNA ISH, immunofluorescence and high content microscopy was performed in iPS-CMs to study the interaction between antisense transcripts, TTN mRNA and splice factor protein RBM20.

**Results:** In mapping *TTN* antisense transcription, we found that *TTN-AS1-276* was the predominant transcript in the human heart and that it was mainly localized in cardiomyocyte nuclear chromatin. Knock down of *TTN-AS1-276* in human iPS-CMs resulted in decreased interaction between the splicing factor RBM20 and *TTN* pre-mRNA, decreased *TTN* I-band exon skipping, and markedly lower expression of the less elastic TTN isoform N2B. The effect on *TTN* exon usage was independent of sense-antisense exon overlap and polymerase II elongation rate. Furthermore, knockdown resulted in longer sarcomeres with preserved alignment, improved fractional shortening and relaxation times.

**Conclusions:** We demonstrate a role for the cardiac *TTN* antisense transcript *TTN-AS1-276* in facilitating alternative splicing of *TTN* and regulating sarcomere properties. This transcript could constitute a target for improving cardiac passive stiffness and diastolic function in conditions such as heart failure with preserved ejection fraction.

## Introduction

Titin (TTN) is the largest protein in the human body (∼3 MDa) and constitutes a central component of the sarcomere.^1^ It is anchored in the Z-disc and extends to the M-band, aligned along the length (∼1μm) of an entire half-sarcomere.^2^ The presence of extensible PEVK-repeat and immunoglobulin-domains in the I-band region gives TTN elastic properties, acting as a biomechanical spring that contributes to sarcomere passive tension and provides elastic force in response to stretch.^3, 4^ In the heart, TTN is a major determinant of myocardial passive stiffness^1, 5^ and plays important roles in regulating diastolic function.^6–8^ Elucidation of molecular mechanisms regulating TTN compliance could thus be of therapeutic importance for diseases characterized by excessive myocardial stiffness and diastolic dysfunction, such as heart failure with preserved ejection fraction (HFpEF).

The mechanical properties of TTN are primarily regulated at the post-transcriptional level, where a series of exon-skipping events among I-band exons produces an array of transcript isoforms differing in the number of extensible domains.^9^ Alternative splicing of the enormous 363 exon *TTN* gene is extraordinarily complex and presumably orchestrated through a highly regulated process involving a number of splicing factors. However, RNA binding motif protein 20 (RBM20) is currently the only protein known to promote alternative splicing of *TTN*.^10^ RBM20 interacts with intronic motifs in pre-mRNA^11^ and prevents inclusion of up- and downstream exons through interference with U1 snRNP splice site recognition.^12^ However, mechanisms regulating recruitment of RBM20 to *TTN* pre-mRNA or additional factors affecting alternative splicing of TTN have not been elucidated.

Antisense transcription, *i.e.* transcription of non-coding RNA from the opposite strand of a coding gene, is pervasive throughout the human genome^13^ and can influence expression and splicing of the sense transcript. Morrissy *et al* found a striking association between the presence of antisense transcription and alternative splicing of the sense gene across the human genome and postulated that slowing of the RNA Polymerase II elongation rate at overlapping sense-antisense exons promotes alternative splicing of the sense transcript.^14^ Others have reported that antisense transcripts can mask splice sites in the sense transcript by forming an RNA-duplex with complementary sequences in the coding transcript^15^ or recruit specific components of the spliceosome to the sense gene pre-mRNA.^16, 17^ The *TTN* locus spans >280 kb on chromosome 2q31 and harbours >80 annotated antisense transcripts, the function of which have not been studied in the heart previously.

The aims of this study were to map *TTN* antisense transcription in the human heart, to elucidate its possible role in regulation of *TTN* splicing, and to study the potential downstream implications of targeting specific *TTN* antisense transcripts on sarcomere organization and function.

## Methods

### Human samples

Left ventricular biopsies from unused organ donor hearts (three women and four men with a mean age of 57 and free from heart disease) were collected at Sahlgrenska University Hospital, Gothenburg, Sweden, embedded in tragacanth and stored at -80°C. The study was approved by the Ethics Board at Gothenburg University.

### Human heart muscle cells

Human iCell iPS-derived cardiomyocytes (iPS-CM) were sourced from FujiFilm Cellular Dynamics Inc. (Madison, WI, USA). Cells were thawed and grown in Plating or Maintenance Medium according to the manufacturer’s instructions. All treatments were carried out five days post plating.

### RNA isolation

25 mg of frozen tissue (see above) was cut into small pieces using a scalpel and transferred to a 2 ml tube. Tissue was homogenized in 700 μl of QIAzol (Qiagen, Hilden, Germany) using an Omni TH rotor-stator homogenizer equipped with disposable Hard Tissue Probes. For isolation of RNA from cells, 700 μl of QIAzol was added directly to cell culture plates after removal of culture medium and washing once with PBS. For isolation of chromatin-enriched and soluble nuclear RNA fractions from cells, the protocol described by Werner et al^18^ was used. Total RNA was isolated using the miRNeasy mini kit (Qiagen) according to the manufacturer’s instructions. The quantity and quality of isolated RNA was assessed with Qubit Flex (ThermoFisher) using the QuantIT RNA HS Assay Kit (ThermoFisher) and Agilent 4200 TapeStation (Agilent Technologies, Santa Clara, CA, USA) using the RNA ScreenTape Analysis Kit (Agilent Technologies).

### siRNA and plasmid DNA transfection

Cells were transfected with Silencer Select siRNA (ThermoFisher, Waltham, MA, USA) towards human *TTN-AS1* (ENST00000659121.1, #n294437), *RBM20* (ENST00000369519.4, #s49081) or a scrambled negative control siRNA sequence (#4390843). For visualization and tracking of sarcomeres, cells were transfected with a plasmid expressing *ACTN2* (*NM_001103.4*) with a eGFP tag.^19^ The plasmid was a gift from Johannes Hell (Addgene plasmid ref #52669). For immunoprecipitation of RBM20, cells were transfected with a pEZ-M03 vector (GeneCopoeia, Rockville, MD, USA) expressing RBM20 (ENST00000369519.4) with an eGFP tag. Transfections were performed using Lipofectamine 3000 (ThermoFisher) according to the manufacturer’s instructions.

### RNA in situ hybridization of human cardiac tissue

10 μm human cardiac cryosections from frozen tissue biopsies (see above) were fixed, dehydrated and pre-treated with Protease IV for the RNAScope Fluorescent Multiplex Assay (Advanced Cell Diagnostics, Hayward, CA, USA) according to the manufacturer’s recommendations (ACD User Manual “320513-USM”). The RNAScope Multiplex Fluorescent Assay was performed using the probe Hs-TTN-C1 (#550361, Advanced Cellular Diagnostics) targeting the ubiquitously expressed exons 24-28 in *TTN* (ENST00000456053.5) and the probe Hs-TTN-AS1-C2 (#1115141-C2, Advanced Cellular Diagnostics) targeting exons 1-5 in *TTN-AS1* (ENST00000456053.5) using the Amp4 Alt C-FL option, according to the manufacturer’s recommendations (ACD User Manual “320293-USM”). Before mounting, 10 μg/ml Wheat Germ Agglutinin-AlexaFluor488 (ThermoFisher) was added to the sections for 10 minutes. Sections were then washed 3x5 min in PBS, counterstained with DAPI for 30 s and mounted using ProLong Gold Antifade Mountant. Sections were imaged using a Nikon TiE TIRF microscope (Nikon Corporation, Tokyo, Japan) equipped with a Photometrics Prime95B sCMOS camera (Teledyne Photometrics, Tucson, AZ, USA).

### Combined RNA in situ hybridization and immunofluorescence

50,000 human iPS-CM were seeded/well in Lab-Tek 4-well chamber slides (Sigma-Aldrich) and cultured for five days according to the manufacturer’s instructions. Cells were then fixed, de- and rehydrated and treated with Protease III in preparation for RNAScope Multiplex Fluorescent Assay (Advanced Cellular Diagnostics) according to the manufacturer’s instructions (ACD Technical Note “320538”). The RNAScope Multiplex Fluorescent Assay was performed according to the manufacturer’s recommendations (ACD User Manual “320293-USM”) with the probes mentioned above. Before mounting, immunofluorescent staining was performed according to the ACD Technical Note “323100-TN”. Slides were blocked with 3% BSA in TBS and stained with a rabbit anti-RBM20 antibody (Abcam, #ab233147) at 10 μg/ml for 1 hour. An AlexaFluor488 anti-rabbit IgG secondary antibody (#4412 Cell Signaling, Danvers, MA, USA) at 1:1000 dilution was then added and incubated for 30 minutes. Slides were counterstained with DAPI for 30 s and mounted using ProLong Gold Antifade Mountant. Slides were imaged using Operetta CLS high content screening instrument (PerkinElmer, Waltham, MA, USA) and the number and localization of *TTN* (Atto 550), *TTN-AS1* (Atto 647) and RBM20 (AlexaFluor488) fluorescent foci were analysed using Harmony 5.2 software.

### Protein gel electrophoresis

Analysis of TTN protein isoforms was performed with agarose gel electrophoresis according to a previously established protocol.^20^ Briefly, human iPS-CM (200,000/well) were seeded in 6-well plates, cultured and transfected with siRNA as described elsewhere. At the end of the experiment, cells were washed once with PBS, scraped in 100 μl of sample buffer (8 M Urea, 2 M Thiourea, 0.05 M Tris-HCl, 75 mM DTT and 3% SDS) and transferred to tubes. Mouse cardiac tissue was frozen in liquid nitrogen, pulverized in a cryogrinder and homogenized in sample buffer using a dounce homogenizer. All samples were heated at 60 °C for 10 minutes, cleared by centrifugation at 13,000x*g* for 5 minutes and stored at -80 °C. A 1.5 mm thick 16x16 cm 1% SeaKem Gold agarose (Lonza group ltd., Basel, Switzerland) gel was cast and used for electrophoresis. 0.5 % 2,2,2-Trichloroethanol (Sigma-Aldrich) was added to the gel to allow for stain-free visualization of proteins. Protein samples were mixed with 4x Laemmli sample buffer supplemented with 0.1 volumes of β-mercapto ethanol before running the gel. ∼20 μl of sample was added per well and electrophoresis was allowed to run at 60V until the dye front ran out of the stacking gel (∼5-6 h). Protein bands were visualized on ChemiDoc MP imaging system (Bio-Rad) using “Stain-free gel” settings. Bands representing TTN isoforms were quantified using densiometric measurements in Image Lab 6.1 (Bio-Rad) and normalized to the myosin heavy chain 7 (MYH7) band.

### RNA Immunoprecipitation

Human iPS-CM were seeded in 60 mm cell culture dishes (800,000 cells/dish) and cultured according to the manufacturer’s instructions. After five days, cells were transfected with pCMV-RBM20-GFP plasmid DNA and siRNA as described elsewhere. 72 hours after transfection, cells were washed twice, scraped in 1 ml of ice-cold PBS and pelleted by centrifugation. 200 μl of mild lysis buffer (Merck) was added and incubated on ice fo 15 minutes. The cell lysate was cleared by centrifugation at 16,000x*g* for 10 minutes. RNA immunoprecipitation was performed on 100 μl cell lysate per sample using the Magna RIP Kit (Merck) according to the manufacturer’s instructions. 5 μg of rabbit polyclonal anti-GFP antibody (#ab290, Abcam) or 1 μg of rabbit IgG antibody was used per immunoprecipitation.

### Chromatin immunoprecipitation

Human iPS-CM were seeded in 10 cm cell culture dishes (2x10^6^ cells/dish) cultured according to the manufacturer’s instructions. Preparation of cross-linked chromatin and chromatin immunoprecipitation was performed using the SimpleChIP Kit (CellSignaling) according to the manufacturer’s instructions. 5 μg of crosslinked chromatin and 1 μg of ChIPAb+ anti- RNA Pol II mouse monoclonal antibody or mouse IgG antibody was used per immunoprecipitation.

### Sarcomere tracking

iPS-CM (∼30,000) were seeded in the middle lower right chamber (Pattern 11) of a U-Slide 8 well^high^ u-Pattern^RGD^ chamber slide (Ibidi GmbH, Grafelfing, Germany). After five days in culture, cells were transfected with siRNA to knock down RBM20 or TTN-AS, and pACTN2- GFP to visualize the sarcomere Z-discs. The procedure for transfection is described in detail above. 48h after transfection, live cell imaging of fluorescently labeled, contracting iPS-CM was performed with wide-field epifluorescence microscopy using a ECLIPSE Ti2-E microscope (Nikon) equipped with a SPECTRA X light engine (Lumencor Inc, Beaverton, OR, USA) and a 60X/1.27 numerical aperture CFI SR Plan Apo objective. Videos of contracting cells, capturing at least two contractions (∼2-3 seconds), were recorded at 30 frames per second using a Nikon DS-Qi2 CMOS camera. Segmentation of z-discs and sarcomere tracking was then performed on a total of 32 video files in the SarcGraph Software.^21^

### qPCR and RT-PCR

cDNA was synthesized using the RevertAid FirstStrand cDNA Synthesis Kit with random hexamer primers (ThermoFisher) according to the manufacturer’s instructions and used in qPCR reactions with 2x Universal TaqMan Master Mix (ThermoFisher) or in RT-PCR reactions with 2x PCR Master Mix (ThermoFisher). The expression of *TTN*, *RBM20*, *GAPDH, CAMK2D, CACNA1C, LMO7, PRKRA, CCDC141, PLEKHA3, FKBP7* and *DFNB59* was assessed with TaqMan Gene Expression Assays (ThermoFisher). For the quantification of specific splice products or exons from *TTN*, *TTN-AS1*, *CAMK2D*, *CACNA1C* and *LMO7*, custom PrimeTime qPCR Probe Assays spanning specific exon-exon junctions or within exons were designed using the PrimerQuest Tool (Integrated DNA Technologies, Coralville, IA, USA). See Supplemental Table for primer and probe sequences. All qPCR reactions were run on a Bio-Rad CFX 96 instrument (Bio-Rad, Hercules, CA, USA). For gene expression analysis, Ct-values were normalized to the reference gene *GAPDH* and expressed relative to the mean of the control group. For quantification of splicing products/isoforms, Ct-values were normalized to a qPCR assay designed to measure all transcripts from the corresponding gene and expressed relative to the mean of the control group. For quantification of immunoprecipitated RNA, Ct-values were transformed to “% of input” using the Ct-value of the 2% Input sample with the formula 2^-(Ct_RIP_-(Ct_Input_-3,32). RT-PCR reactions were run on a 2% agarose gel with GelRed and amplicons were visualized using a ChemiDoc MP imaging system with the “UV Trans” application. Bands corresponding to the PCR amplicons were quantified using densiometric measurements in Image Lab 6.1 and normalized to the input samples.

### RNA-sequencing

For human cardiac tissue samples, 800 ng of RNA was used as input for library preparation using the TruSeq Stranded Total RNA Library Prep Gold kit (Illumina, San Diego, CA, USA) with rRNA Removal Mix. Sequencing was performed on a NovaSeq 6000 system using the NovaSeq 6000 S4 reagent kit (Illumina) with 101 bp paired-end reads.

Raw fastq files were processed using the nf-core RNAseq/3.12.0 pipeline.^22, 23^ Trimming of adapter sequences and removal of low-quality reads were done using Trim Galore 0.6.7^24^ and rRNA was removed by SortMeRNA 4.3.4^25^ with rRNA database smr_v4.3_default_db.fasta. BBmap 38.61b^26^ was used to map the clean reads to the human reference genome (Homo_sapiens.GRCh38.dna_sm.primary_assembly.fa.gz). Some samples were processed using the BBmap function repair.sh before being fed to BBmap for mapping. The generated mapped bam files were sorted and indexed with samtools 1.17.^27^ Mapping quality was evaluated with samtools 1.17 (stats, idxstats and flagstat) and rseqc 2.6.4 (infer_experiment.py, junction_saturation.py, bam_stat.py, read_duplication.py, read_distribution.py and junction_annotation.py).^28^ The biotype content of the reads was counted with the featureCount program from the subread 2.0.3 package.^29^ The unique mapping rate was also estimated by counting the reads with NH:i:1 using samtools 1.17.

For human iPS-derived cardiomyocytes, 10 ng of RNA was used as input for cDNA synthesis using the SMART-Seq v4 Ultra Low Input RNA Kit for Sequencing (Takara Bio, San Jose, CA, USA). Library preparation was performed using the Nextera XT DNA Library Preparation Kit (Illumina). Sequencing was performed using the NovaSeq 6000 SP Reagent Kit v 1.0 (Illumina) with 101 bp paired-end reads.

Demultiplexing was performed using the bcl2fastq2 software (RRID:SCR_015058) with default settings. Reads were aligned to the human GRCh38 reference genome with the gencode version 33 annotation, using the STAR software.^30^ Quantification of gene expression was performed using the featureCounts software. Raw gene counts were transformed to fragments per kilobase of exon per million mapped fragments (FPKM) using an in-house python script, fCounts2fpkm.py.

All mapped reads were then counted with DEXseq 1.44.0^31, 32^ following the instructions at https://rdrr.io/bioc/DEXSeq/f/vignettes/DEXSeq.Rmd. Only reads with mapping quality ≥10 were counted. The counting used the feature annotation information from Homo_sapiens.GRCh38.109.gtf.gz (https://www.gencodegenes.org/human/release_33.html). Percent spliced in (PSI) was estimated using the Calculate-PSI package (https://github.com/jalwillcox/Calculate-PSI).

## Results

### Cardiac antisense transcription in the TTN locus

Antisense transcription in the *TTN* gene locus is extensive, with 81 *TTN-AS1* transcripts annotated in GENCODE v. 44 (**Supplemental Figure 1a**). To obtain an overview of *TTN-AS1* transcript structure and expression in the human heart, we determined exon usage across all *TTN-AS1* isoforms by applying the DEXSeq analysis pipeline on RNA-sequencing data from left ventricular tissue without heart disease (n=7, **Figure 1a**). Exon usage varied substantially within and between transcripts, but the high-expression (75^th^ percentile) exons all belonged to a group of core TTN-AS1 transcripts (**Figure 1b**). We quantified the four TTN-AS1 transcripts with the highest expression exons (TTN-AS1-276, -209, -203 and -223) using the same human cardiac tissue without heart disease (n=7, SHS controls) and custom qPCR assays targeting transcript-specific exons or exon-exon junctions (**Figure 1c**). The highest expression in cardiac tissue was observed for TTN-AS1-276 (ENST00000659121), which is also the ENSEMBL canonical transcript, defined as the single most representative and best-supported transcript from any given gene. TTN-AS1-276 is a 6297 bp transcript with 13 exons spanning >250 kb of the *TTN* gene (**Supplemental Figure 1a**). The transcriptional start site of *TTN-AS1*-276 lies ∼2500 bp downstream of *TTN* and ENCODE ChIP-Seq data shows that this genomic region is characterized by H3K4Me3-enrichment, characteristic for active promoters (**Supplemental Figure 1b**). RNA in situ hybridization (ISH) revealed widespread expression of *TTN-AS1-276* in human left ventricular tissue (**Figure 1d** and **Supplemental Figure 2**), localized predominantly in cardiomyocytes (as defined by large cell/nuclear size and presence of *TTN* ISH foci). Cardiomyocyte-specific expression of *TTN-AS1* was corroborated by GTEx cardiac single nucleus RNA-sequencing data,^33^ where median log-normalized and scaled expression counts for *TTN-AS1* was >10,000-fold higher in cardiomyocytes than in any other cardiac cell type.

**Figure 1.**
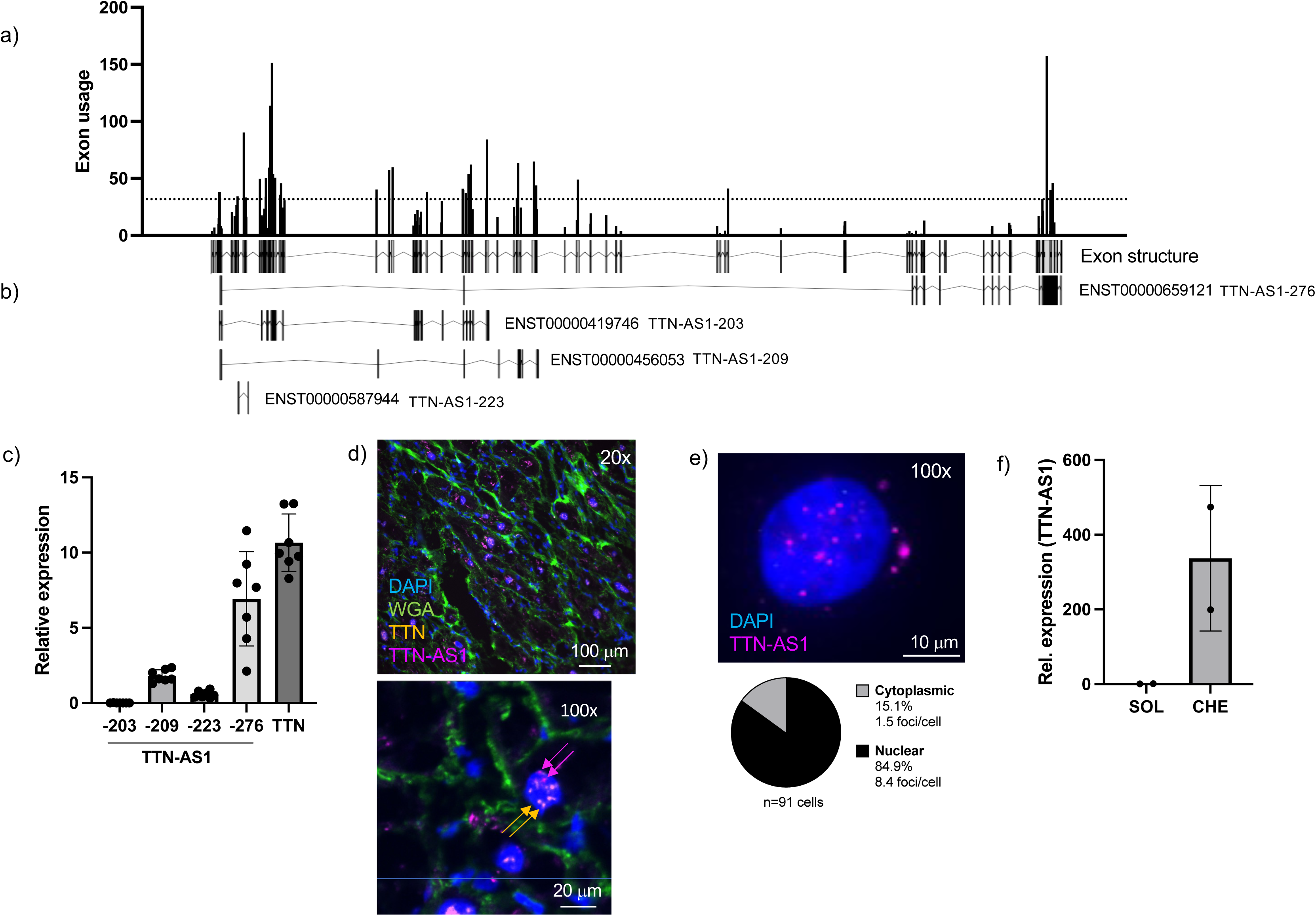
Expression and localization of TTN antisense transcripts in the human heart. A) *TTN-AS1* exon usage was calculated using cardiac RNA-sequencing data from 7 organ donor hearts without heart disease. The dashed line indicates the 75^th^ percentile. B) The *TTN-AS1* transcripts with the highest exon usage. C) Relative expression of *TTN-AS1* transcripts and *TTN* with qRT-PCR in the 7 organ donor hearts without heart disease. D) RNA in situ hybridization (ISH) for *TTN-AS1* (magenta) and *TTN* (yellow) in a tissue section from one representative human cardiac biopsy. Cell membranes were stained with wheat germ agglutinin (WGA). E) RNA ISH for *TTN-AS1* in human iPS-derived cardiomyocytes (iPS-CM). Nuclei were stained with DAPI. Below, quantification of the proportion of nuclear and cytoplasmic *TTN-AS1* ISH foci. F) Relative expression of TTN-AS1 in iPS-CM nuclear compartments with qRT-PCR. Data are derived from two different RNA preparations per group. SOL: soluble fraction, CHE: chromatin-enriched fraction.

Clues regarding the functional role of a natural antisense transcript can be drawn from its intracellular localization.^34^ We therefore performed RNA ISH on human iPS-derived cardiomyocytes (iPS-CM) and observed that ∼85% of *TTN-AS1-276* foci were localized in the nucleus (**Figure 1e**). Moreover, fractionation of iPS-CM nuclear RNA revealed ∼300-fold higher levels of *TTN-AS1-276* in the chromatin compartment over the soluble compartment (**Figure 1f**). We conclude that *TTN-AS1-276* is the predominant *TTN* antisense transcript in the human heart and that it is localized to cardiomyocyte nuclear chromatin.

### TTN-AS1 does not regulate TTN expression

Natural antisense transcripts often play a role as cis-acting transcriptional regulators of the sense protein-coding gene.^35^ To assess whether *TTN-AS1-276* affects the expression of *TTN*, we transfected human iPS-CM with siRNA directed towards Exon 1 of *TTN-AS1-276* (siTTN-AS1). A statistically significant, ∼40 % knock down of *TTN-AS1-276* was observed 48 hours after transfection, but no effect on expression of *TTN* or any of the other protein-coding genes immediately within 100 kb up- and downstream of *TTN* (**Supplemental Figure 3**) could be detected. We conclude that *TTN-AS1* does not play a role as a cis-acting transcriptional regulator.

### TTN-AS1 regulates splicing of TTN I-band exons

We next hypothesized that *TTN-AS1* could influence alternative splicing of *TTN*, as the presence of antisense transcription across the human genome has been shown to associate strongly with alternative splicing of the corresponding sense gene.^14^ We performed RNA-seq on human iPS-CM transfected with siTTN-AS1 or, as a positive control, siRNA towards *RBM20* (siRBM20), a splicing factor which mediates exon skipping in TTN^11^, and calculated proportion spliced-in (PSI) for each *TTN* exon. Successful knock down of *TTN-AS1* and *RBM20* was confirmed with qPCR (**Supplemental Figure 4a**). *TTN* PSI in control iPS-CM correlated well (Pearson r=0.837, p<0.0001) with previously reported data from human left ventricular tissue,^36^ with extensive splicing out of exons across the I-band region throughout exons 50-242 (numbered according to the complete inferred TTN meta-transcript**, Supplemental Figure 4b, Figure 2a**). Knock down of *TTN-AS1* caused significantly increased PSI for 26 exons (indicated by red bars in **Figure 2b**). Affected exons were all situated in the I-band and had PSI values in control cells in the range of 70-85%. The most pronounced effect of TTN-AS1 knock down was observed in exons immediately downstream of exon 49 (exons 50-89, indicated by a dashed square in **Figure 2b**). These exons are normally spliced out of *TTN* through RBM20-mediated exon skipping^37^ and alternative splicing/exon skipping from exon 49 produces a range of *TTN* splice products^38^, notable examples of which are shown in **Figure 2c**. Knock down of *RBM20* resulted in a similar increase in PSI in exons downstream of exon 49 (**Supplemental Figure 4c**), confirming its role in mediating I-band exon skipping. To study whether the increased PSI in exons downstream of 49 following TTN-AS1 knock down was caused by decreased exon skipping, we quantified the primary splice products across exon 49 splice junctions in iPS-CM transfected with siTTN-AS1 or siRBM20 using qPCR. We observed a significant decrease in expression of *TTN* splice products dependent on exon skipping and a reciprocal increase in the isoform where exon 49 is spliced directly with exon 50 (**Figure 2d**). We conclude that TTN-AS1 regulates exon usage in I-band *TTN* through facilitating exon skipping downstream of exon 49.

**Figure 2.**
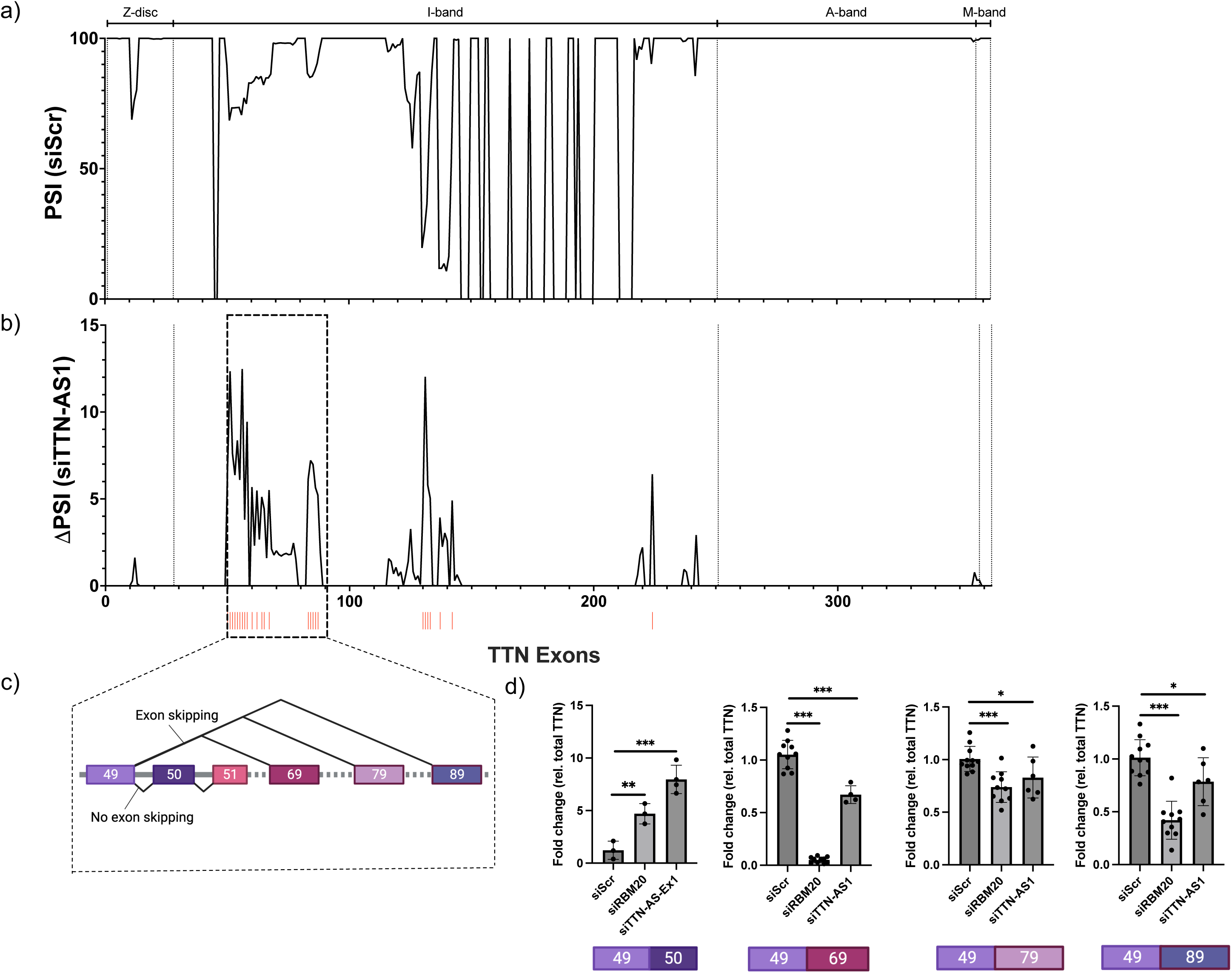
TTN-AS1 regulates inclusion of I-band exons in *TTN*. A) Percent spliced in (PSI) for all exons across the *TTN* gene calculated based on RNA-sequencing reads from human iPS-derived cardiomyocytes (IPS-CM). The line represents the mean of three replicates (individual RNA preparations) per experimental group. B) The difference in PSI (1′PSI) comparing cells transfected with siRNA to *TTN-AS1* (siTTN-AS1) with cells transfected with scrambled negative control siRNA (siScr). Exons with statistically significant 1′PSI are marked with red bars (adjusted p<0.05). C) Depiction of exon skipping from *TTN* exon 49. D) Quantification of *TTN* splice products in iPS-CM transfected with siTTN-AS1, siRBM20 or siScr, using custom qRT-PCR assays spanning the indicated exon-exon junctions. Expression data is normalized to that of total *TTN* and expressed relative to the mean of the negative control cells (siScr). Data is derived from two separate experiments with 3-6 replicates (individual RNA preparations) per experimental group. Differences between each individual experimental group and the control group were assessed with Student’s t-tests, *p<0.05, **p<0.01, ***p<0.001.

### TTN-AS1 regulates TTN isoform composition

Cardiac *TTN* is composed of two primary isoforms, N2BA and N2B, differing in the extent to which I-band domains are included. N2B is produced by splicing of exon 49 with exon 219, excluding many extensible I-band domains and resulting in a short and less elastic TTN protein isoform (**Figure 3a**). As our data showed that *TTN-AS1* promotes exon skipping downstream of exon 49, we hypothesized that knock down of *TTN-AS1* would result in decreased expression of the N2B isoform. To address this, we quantified expression of N2B and N2BA in iPS-CM transfected with siTTN-AS1 or siRBM20 using custom qPCR assays spanning the exon 49-219 junction (N2B) and the exon 108-109 junction (constitutively included in N2BA). We observed significantly lower expression of the N2B isoform in iPS-CM where *TTN-AS1* had been knocked down, whereas expression of N2BA was unaffected (**Figure 3b**). This resulted in a 2-fold increase in the N2BA:N2B ratio (**Figure 3c**). Knock down of *RBM20* caused a dramatic decrease in N2B expression and a pronounced increase in the N2BA:N2B ratio, in line with previous reports.^10, 11^ Next, we analysed the consequence of TTN-AS1 knock down on N2B and N2BA protein isoforms using gel electrophoresis. As previously reported,^36^ iPS-CM expressed a longer, fetal-like N2BA isoform and N2B expression was considerably lower than in adult cardiac tissue (**Figure 3d**). Nevertheless, in line with our observation on the mRNA level, we found N2B protein expression to be significantly lower in iPS-CM where TTN-AS1 had been knocked down but saw no effect on N2BA expression. This was also reflected in a significantly increased N2BA:N2B ratio. Again, the effect of *RBM20* knock down had a similar effect on N2B expression and the N2BA:N2B ratio (**Figure 3f**). The Cronos TTN isoform, which is transcribed from an internal promoter in the distal I-band region (between exon 239 and 240 of the inferred meta transcript), was not affected by either siTTN-AS1 or siRBM20 (**Figure 3g**). We conclude that TTN-AS1 regulates *TTN* isoform composition and can be targeted to promote translation of longer and more extensible TTN.

**Figure 3.**
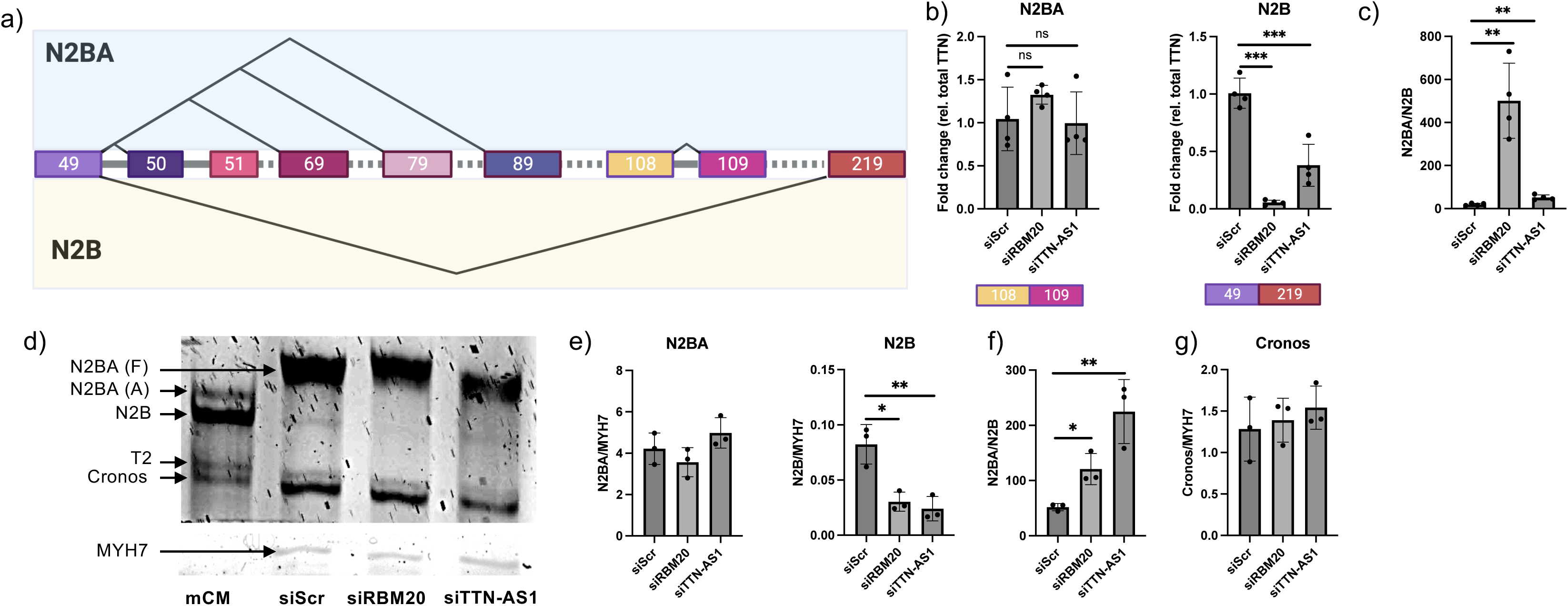
*TTN-AS1* knock down causes a shift in TTN isoform composition. A) Depiction of alternative splicing pathways from *TTN* exon 49 producing the two major cardiac isoforms, N2BA and N2B. B) Quantification of N2BA and N2B isoforms in iPS-CM transfected with siTTN-AS1, siRBM20 or siScr, using custom qRT-PCR assays spanning the indicated exon-exon junctions. Expression data is normalized to that of total *TTN* and expressed relative to the mean of the negative control cells (siScr). Data is derived from two separate experiments with two replicates (individual RNA preparations) per experimental group, ***p<0.001. C) The N2BA to N2B ratio in iPS-CM following transfection siTTN-AS1, siRBM20 or siScr, derived from data in B). D) Gel electrophoresis of cardiac protein from mouse (mCM), or iPS-CM transfected with siScr, siRBM20 or siTTN-AS1. The major TTN isoforms (N2BA, N2B and Cronos) are indicated. T2: TTN degradation product. The band corresponding to MYH7 is shown in the bottom. E, G) Quantification of band intensity for N2BA, N2B and Cronos relative to MYH7. F) Ratio of N2BA to N2B. Data is derived from three separate experiments. Differences between each individual experimental group and the control group were assessed with Student’s t-tests *p<0.05, **p<0.01.

### TTN-AS1 regulates sarcomere function and cardiomyocyte contraction dynamics

TTN isoform composition is a key determinant of cardiomyocyte mechanical properties. As knock down of *TTN-AS1* resulted in increased inclusion of I-band exons and a shift towards longer and more elastic TTN, we wanted to investigate the consequences of *TTN-AS1* knock down on sarcomere structure and dynamics. To this end, we applied sarcomere tracking analysis on live iPS-CM after siRNA-mediated knock down of *TTN-AS1* or *RBM20*. Cells were transfected with a plasmid encoding GFP-tagged alpha-actinin-2 (pACTN2-GFP) to visualize sarcomere Z-discs (**Figure 4a**) and recorded during multiple contractions with live cell imaging (**Supplemental Video 1**). SarcGraph v.0.2.1 was used to detect, track, and analyse functional parameters of 14,636 sarcomeres from >200 cells during contraction (**Figure 4b-h**). Results showed that sarcomere length was increased by ∼10% (p<0.001) in cells where *TTN-AS1* had been knocked down (**Figure 4e**), whereas the sarcomere alignment, as measured by the orientational order parameter (OOP), was unaffected **(Supplemental Figure 5**). Expectedly, knock down of RBM20 caused a similar increase in sarcomere length (p<0.001). Fractional shortening (FS) was increased from 7.5% in control cells to 9% in cells transfected with siTTN-AS1 (p<0.001), indicating increased contractility (**Figure 4f**) and both mean contraction (**Figure 4g**) and mean relaxation (**Figure 4h**) time increased by ∼30-40% (p<0.001) following TTN-AS1 knock down. Knock down of RBM20 mirrored these effects as well. We conclude that knock down of TTN-AS1 results in longer and more compliant sarcomeres with improved contractile properties.

**Figure 4.**
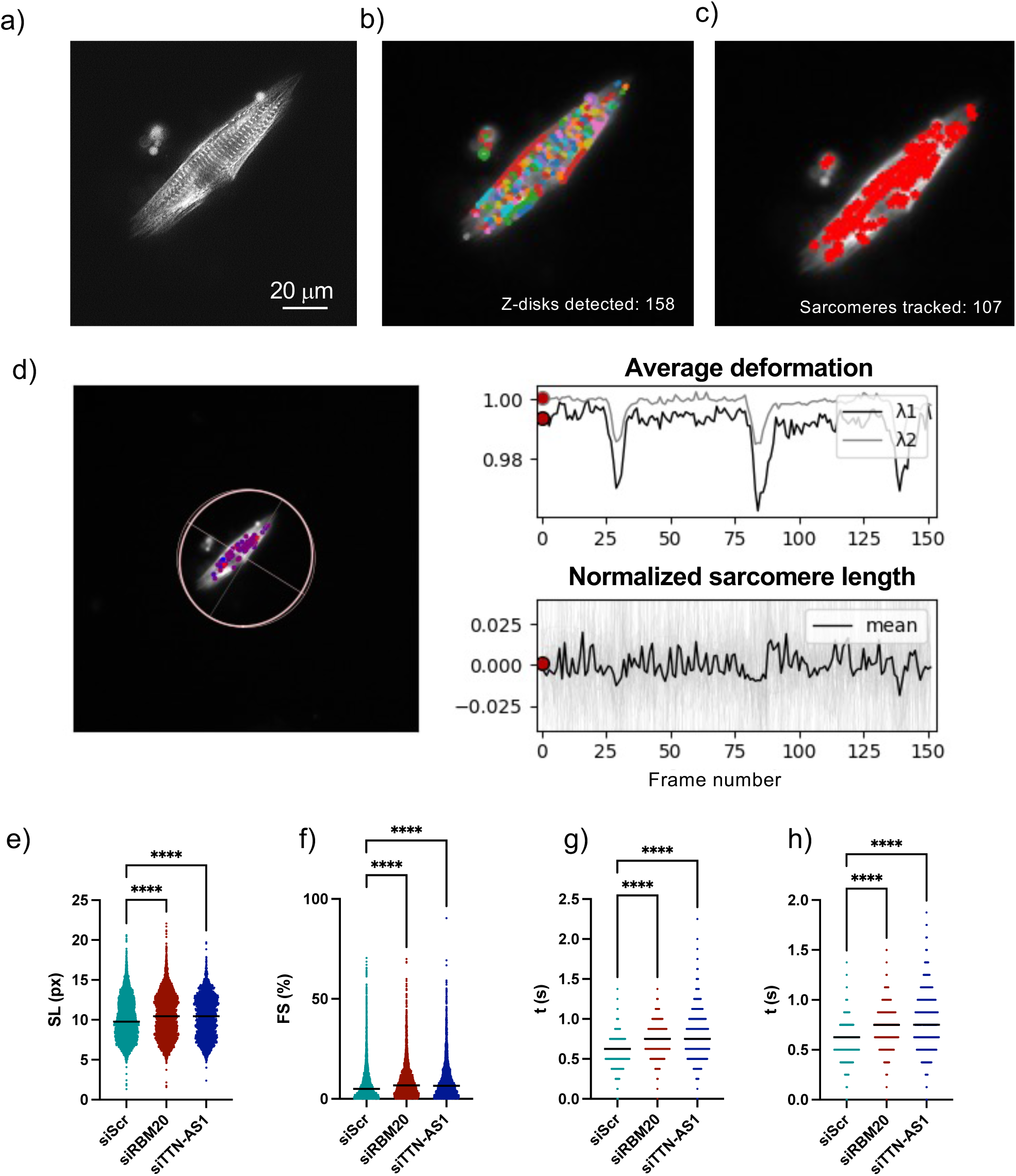
*TTN-AS1* knock down results in increased sarcomere length and altered contraction dynamics. A) Representative fluorescent image of a human iPS-derived cardiomyocyte transfected with a ACTN2-GFP plasmid for visualization and tracking of sarcomeres in SarcGraph software. Visualization of B) Z-disc and C) sarcomere segmentation in SarcGraph software. D) Time series data for average deformation and normalized sarcomere length during three contractions. E) Sarcomere length (SL), F) fractional shortening (FS), G) contraction time and H) relaxation time from a total of 14,636 segmented iPS-CM sarcomeres following transfection with siTTN-AS1, siRBM20 or siScr. Data are derived from two separate experiments. Differences between each individual experimental group and the control group were assessed with Student’s t-tests, ****p<0.0001.

### No impact of overlapping antisense transcript exons on TTN exon usage and Pol II elongation rate

Next, we wanted to explore the mechanism by which TTN-AS1 affects alternative splicing of TTN. Morrissy *et al* hypothesized that a reason for the increased alternative splicing observed in exons with overlapping antisense exons is that polymerase elongation rates are decreased,^14^ as slower RNA polymerase II (Pol II) rates have been shown to increase the rate of alternative splicing.^39^ When considering exons from all annotated *TTN-AS1* and *TTN* transcript isoforms in GENCODE, ∼16% of *TTN* exons overlap with one or more *TTN-AS1* exons (red bars in **Figure 5a**). Apart from the Z-disc, overlapping exons are present throughout the whole *TTN* gene. If the presence of an antisense exon was to promote exon skipping in *TTN*, we expected usage of such *TTN* exons to be lower compared to those without an antisense exon. To analyse this, we leveraged DEXSeq exon usage data from human left ventricular tissue without heart disease (n=7), but the results revealed instead a significantly increased usage of TTN exons with an overlapping antisense exon (**Figure 5a-b**), contradicting this hypothesis. Moreover, we observed no enrichment of exons with an overlapping antisense exon among the 26 *TTN* exons with increased PSI following *TTN-AS1* knock down (12% vs 16% in *TTN* overall).

**Figure 5.**
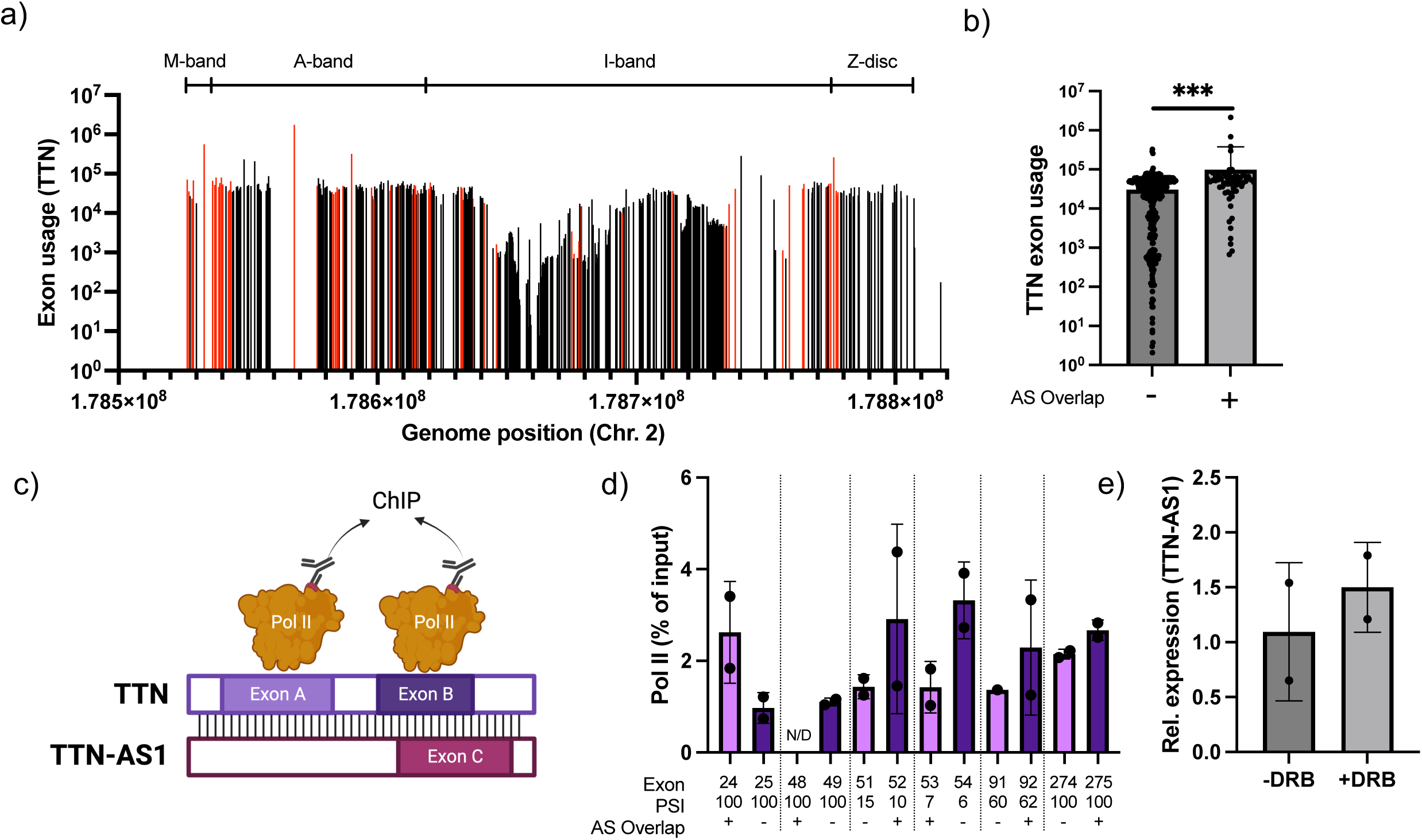
No influence of overlapping antisense exons on TTN exon usage. A) Mean *TTN* exon usage was calculated using cardiac RNA-sequencing data from 7 organ donor hearts without heart disease. Exons with an overlapping *TTN-AS1* exon are marked in red. B) Usage of exons with and without an overlapping TTN-AS1 exon. The difference in exon usage between exons with and without an overlapping antisense exon was assessed with Student’s t-test, ***p<0.001. C) Depiction of experimental design for PolII ChIP experiments in D). ChIP was performed with a PolII antibody on chromatin from human iPS-derived cardiomyocytes (iPS-CM). PolII occupancy was assessed on six pairs of consecutive *TTN* exons where one had an overlapping antisense exon and the other did not, using qRT-PCR on ChIP DNA. Data was derived from two separate experiments. E) Relative expression of *TTN-AS1* in the chromatin-enriched fraction of iPS-CM nuclei after treatment with DRB, analyzed with qRT-PCR. Data was derived from two separate experiments.

We next investigated Pol II occupancy across a selection of *TTN* exons with and without *TTN-AS1* exon overlap in iPS-CM using ChIP-PCR. We designed primers for 6 pairs of adjacent *TTN* exons where one exon had an overlapping antisense exon and the other did not (**Figure 5c**), across different TTN domains and reflecting varying PSIs (**Figure 5d**). With the exception of exon 48, for which no Pol II signal could be detected, there was no robust or meaningful difference in Pol II occupancy at exons with overlapping antisense exons compared to adjacent exons without overlapping antisense exons (**Figure 5d**). Moreover, inhibition of Pol II elongation by addition of DRB did not affect the degree of *TTN-AS1* chromatin enrichment in iPS-CM, indicating that Pol II pausing does not influence *TTN-AS1* tethering to chromatin (**Figure 5e**). Taken together, these data show that *TTN-AS1* influences *TTN* exon usage/alternative splicing in a manner independent of specific sense-antisense exon overlap and Pol II elongation rate.

### TTN-AS1 facilitates interaction between RBM20 and TTN mRNA

We then wanted to explore alternative mechanisms whereby *TTN-AS1* could regulate alternative splicing of *TTN*. Antisense transcripts have previously been reported to influence splicing either by forming an RNA-duplex with complementary sequences in the coding transcript, thereby masking splice-sites in pre-mRNA of the sense gene^15^ or by recruiting or guiding specific components of the spliceosome to the sense gene pre-mRNA.^16, 17^ We reasoned that TTN-AS1 could exert its function through either of these mechanisms, but given the similar patterns in *TTN* ΔPSI, *TTN* isoform composition and functional consequences on sarcomere dynamics in iPS-CM following knock down of *TTN-AS1* and *RBM20*, we believed that a mechanism whereby *TTN-AS1* recruits RBM20 to *TTN* mRNA would be more plausible. To explore these hypotheses, we analysed the quantity and nuclear localization of *TTN-AS1*, *TTN* mRNA and RBM20 protein in iPS-CM using combined RNA ISH and immunofluorescence. First, we found that nuclear *TTN*:*TTN-AS1* co-localization, indicative of duplex formation, was rare, occurring once in every fifth cell on average, and interestingly, was then almost exclusively (>90% of instances) observed as part of a cluster involving RBM20 (**Figure 6a** and **Supplemental Figure 6a**), with a fluorescence pattern indicative of RBM20-mediated *TTN* splicing.^37, 40^ This observation contradicts a mechanism involving direct TTN:TTN-AS1 duplex formation and strengthens the hypothesis that *TTN-AS1* regulates *TTN* splicing via RBM20. To test this hypothesis further, we quantified the proportion of *TTN* RNA ISH foci co-localized with RBM20 in iPS-CM transfected with siTTN-AS1 using high content imaging analysis. In control cells, we estimated that ∼7.5% of all nuclear *TTN* RNA ISH foci co-localized with RBM20 (**Figure 6b**). Interestingly, in iPS-CM where *TTN-AS1* had been knocked down, the proportion of *TTN* foci co-localized with RBM20 was more than halved (p<0.0001, **Figure 6c**). This suggests that *TTN-AS1* facilitates interaction between RBM20 and TTN pre-mRNA. We then sought to corroborate these findings using RNA immunoprecipitation (RIP) on iPS-CM. Since there are no commercially available RIP-grade antibodies for RBM20, we instead transfected iPS-CM with a plasmid encoding a RBM20-GFP fusion protein (pRBM20-GFP) and used a GFP antibody to pull down the RBM20 RNA interactome (**Figure 6d**). Transfection with pRBM20-GFP resulted in transgenic expression of RBM20 fusion protein (**Supplemental Figure 7a**). We observed significant enrichment of both *TTN* and *TTN-AS1* in RBM20-RIP RNA compared to unrelated *GAPDH* RNA (**Figure 6e**) and to negative control IgG IP (**Supplemental figure 7b**). Expectedly, the *TTN-AS1* RIP signal was significantly reduced in cells where *TTN-AS1* had been knocked down. Interestingly, there was also significantly less *TTN* interacting with RBM20 after *TTN-AS1* knock down, giving additional support for a mechanism whereby *TTN-AS1* facilitates interaction between RBM20 and *TTN* mRNA (**Figure 6e**).

**Figure 6.**
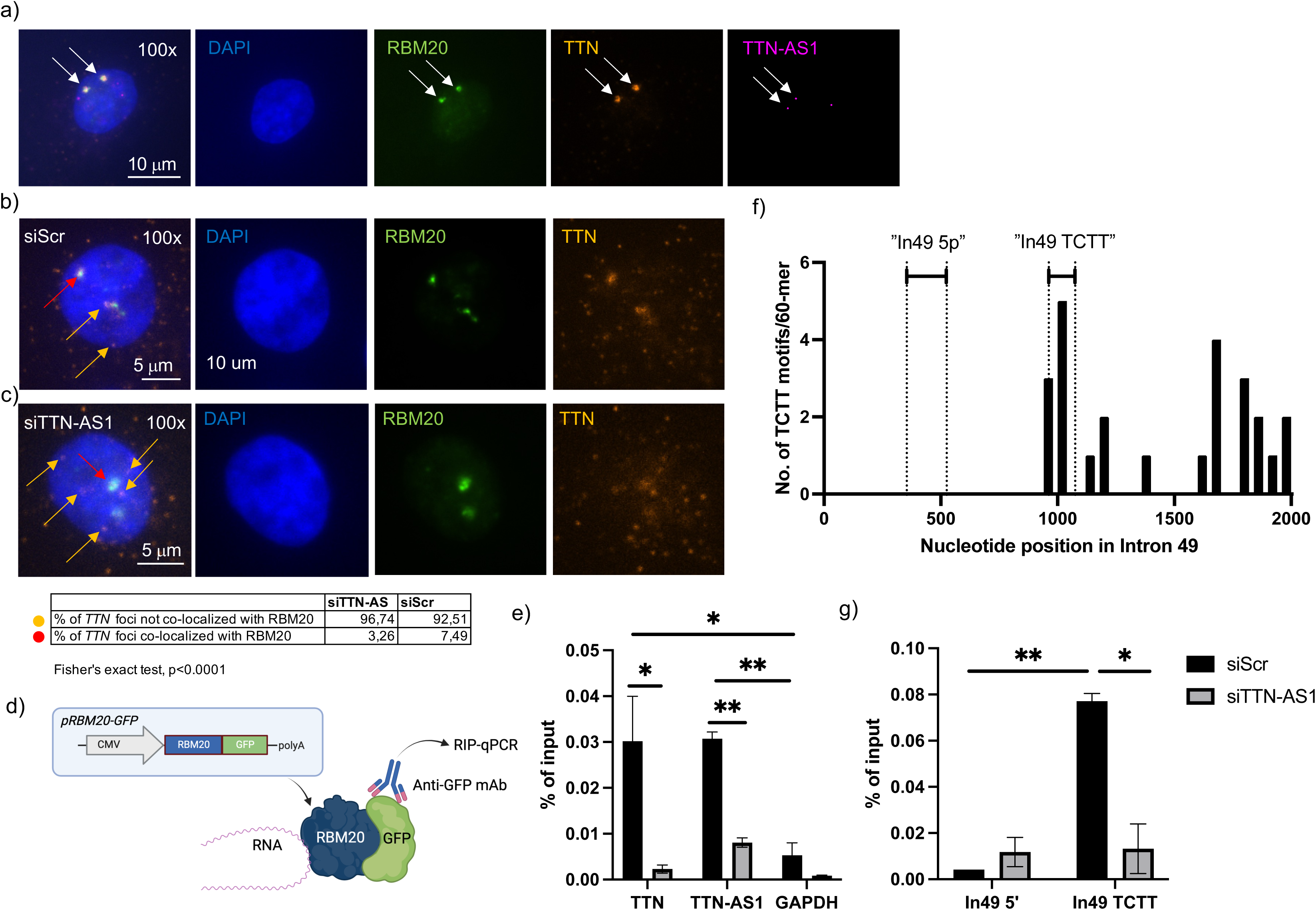
TTN-AS1 facilitates interaction between RBM20 and TTN mRNA. A-C) Human iPS-derived cardiomyocytes (iPS-CM) were subjected to combined RNA in situ hybridization (ISH) for *TTN-AS1* (magenta) and *TTN* (orange) and immunofluorescence for RBM20 (green) following transfection with siRNA towards *TTN-AS1* (siTTN-AS1), *RBM20* (siRBM20) or scrambled negative control siRNA (siScr). Nuclei were counterstained with DAPI. Co-localization of TTN and RBM20 foci were analyzed with high content imaging. D) Depiction of experimental design for the RBM20 RNA immunoprecipitation (RIP) experiment. iPS-CM were transfected with a plasmid expressing a RBM20-GFP fusion protein. RIP was performed on iPS-CM protein using a GFP antibody and qRT-PCR was used to analyze immunoprecipitated RNA. E) Enrichment of *TTN* and *TTN-AS1* in GFP-RBM20 RIP RNA from iPS-CM transfected with siTTN-AS1 or siScr, analyzed with qRT-PCR. Analysis of unrelated *GAPDH* RNA was included as a negative control. Data are derived from two separate experiments with two technical replicates (individual immunoprecipitates) in each group *p<0.05, **p<0.01. F) The number of TCTT motifs/60 bp across intron 49 of *TTN*. The positions of RIP-qPCR assays used in G) is indicated with dashed lines. G) qPCR of GFP-RBM20 RIP RNA from E) using assays targeting regions in intron 49 without TCTT-motifs (“In49 5p”) and enriched with TCTT-motifs (“In49 TCTT”). Data are derived from two separate experiments with two technical replicates (individual immunoprecipitates) in each group. Differences in the RIP-signal between cells transfected within and between groups were assessed using ANOVA with Dunnett’s multiple comparisons test, *p<0.05, **p<0.01.

RBM20 mediates exon skipping by interacting with intronic TCTT motifs in pre-mRNA.^11^ We hypothesized that *TTN-AS1* assists in the binding of RBM20 to such intronic motifs upstream of skipped exons. To test this hypothesis, we chose to focus on intron 49, with which RBM20 would interact for skipping of exon 50 to occur. We scanned the sequence of intron 49 and found an enrichment of TCTT-motifs in the middle of the intron and towards the 3’ end, whereas the 5’ end of the intron was completely devoid of TCTT-motifs (**Figure 6f**). To assess binding of RBM20 to different regions of intron 49, we designed qPCR assays targeting a 5’ region free from TCTT-motifs (“In49 5p”) and a region enriched in TCTT-motifs in the middle of the intron (“In49 TCTT”). We then performed RBM20-RIP qPCR using these assays and expectedly, found that RBM20 was significantly enriched in the TCTT-rich region (**Figure 6g**). Interestingly, RBM20 binding to the TCTT-rich region was significantly reduced upon knock down of *TTN-AS1* (**Figure 6g**). Taken together, these results point towards a mechanism whereby *TTN-AS1* mediates exon skipping in *TTN* through interacting with and recruiting RBM20 to intronic TCTT motifs.

### TTN-AS1 mediates splicing of other RBM20 targets

In a previous study, Bertero et al postulated that RBM20 forms a *trans-*interacting chromatin domain (TID), driving spatial proximity of multiple genomic loci representing a network of different RBM20 targets.^40^ The assembly of this splicing complex is initiated at the *TTN* genomic locus, where transcription of *TTN* nucleates RBM20 foci, which drives formation of the TID. Given the overlap and spatial proximity of *TTN-AS1* and *TTN* loci, the evidence of physical interaction of *TTN-AS1* and *TTN* RNA with RBM20 protein and the observation that the number of RBM20 protein foci decreased in iPS-CM after TTN-AS1 knock down (**Supplemental Figure 6**), we hypothesized that *TTN-AS1* might facilitate the assembly of the RBM20 splicing factory. If this was the case, knock down of *TTN-AS1* would affect splicing of not just *TTN*, but also of other RBM20 targets. To explore this hypothesis, we first analyzed PSI data from iPS-CM where either *TTN-AS1* or *RBM20* had been knocked down, focusing on three genes shown to be included in the RBM20 TID: *CACNA1C*, *CAMK2D* and *LMO7* (**Supplemental figure 8a**). In line with previous reports, there was significantly different 1′PSI for Exon 8 and 30 in *CACNA1C* (**Figure 7a**), Exon 14 in *CAMK2D* (**Figure 7b**), and Exon 9-11 in LMO7 (**Figure 7c**) following RBM20 knock down. Interestingly, this effect was mirrored almost exactly in cells where *TTN-AS1* had been knocked down. Guided by these results, we quantified splicing products using qPCR assays spanning isoform-specific exon-exon junctions (*CACNA1C* and *LMO7*) or with RT-PCR (*CAMK2D*). For *CACNA1C*, we saw a significant downregulation of the alternative exon 8a relative to exon 8 in both cells transfected with siRBM20 and siTTN-AS1 (**Figure 7d**). We also observed significant downregulation of LMO7 isoforms subject to exon 9 and 10 skipping following knock down of both siRBM20 and siTTN-AS1 (**Figure 7e**). Finally, we detected a significant shift from CAMK2D-C, which requires skipping of exon 14-16, to the CAMK2D-B isoform, which requires skipping of exon 15, and unspliced CAMK2D, in both experimental groups (**Figure 7f** and **Supplemental figure 8b**). Results were overall strikingly similar between siRBM20 and siTTN-AS1, with the only exception of the CAMK2D-9 isoform, where knock down of RBM20 caused a significant decrease, and knock down of TTN-AS1 instead caused a significant increase. Next, we confirmed the interaction between RBM20 protein and *CACNA1C*, *LMO7* and *CAMK2D* RNA by RIP-qPCR (**Figure 7g**). All three genes showed significant enrichment compared to negative control RNA (*GAPDH*) in RBM20-RIP RNA and compared to the signal from non-specific IgG RIP (**Supplemental Figure 8c**). As expected, the RIP signals for all three genes were almost completely abolished upon TTN-AS1 knock down (**Figure 7g**). In all, these results support a mechanism where *TTN-AS1* is a component of the RBM20 splicing machinery and abets alternative splicing of RBM20 target genes in cardiomyocytes.

**Figure 7.**
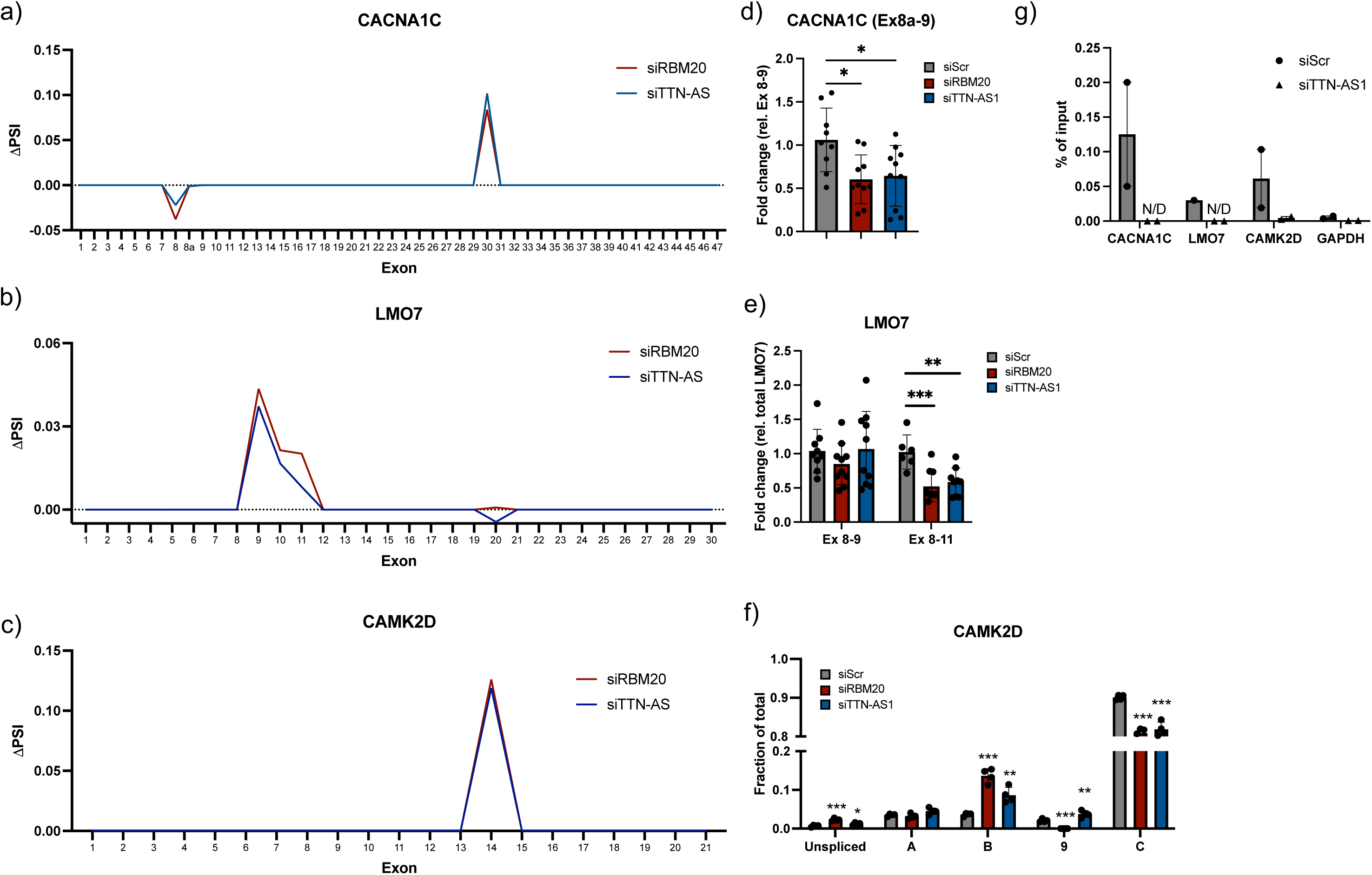
TTN-AS1 facilitates splicing of additional RBM20 targets. A-C) Difference in PSI (1′PSI) comparing cells transfected with siRNA to *TTN-AS1* (siTTN-AS1, blue) or *RBM20* (siRBM20, red) with cells transfected with scrambled negative control siRNA (siScr) across all exons of the A) *CACNA1C,* B) *LMO7* and C) *CAMK2D* genes in human iPS-derived cardiomyocytes (iPS-CM). The lines represent the mean of three technical replicates (individual RNA preparations). D-F) Relative expression of alternative splice products of the D) *CACNA1C*, E) *LMO7* and F) *CAMK2D* genes in iPS-CM transfected with siRBM20, siTTN-AS1 or siScr and analyzed with qRT-PCR (*CACNA1C* and *LMO7*) and semi-quantitative RT-PCR (*CAMK2D)*, respectively. Data is derived from three separate experiments with three technical replicates (individual RNA preparations) in each group. Differences between each individual experimental group and the control group were assessed with Student’s t-tests,*p<0.05, **p<0.01, ***p<0.001. G) Enrichment of *CACNA1C, LMO7* and *CAMK2D* in GFP-RBM20 RIP RNA from iPS-CM transfected with siTTN-AS1 or siScr, analyzed with qRT-PCR. Analysis of unrelated *GAPDH* RNA was included as a negative control. Data are derived from two separate experiments.

## Discussion

In this study, we comprehensively map *TTN* antisense transcription in human heart tissue and define a functional role for the most abundant transcript, TTN-AS1-276, in alternative splicing of *TTN*. Based on several lines of evidence from different techniques, we postulate that *TTN* antisense transcripts interact with RBM20 to facilitate exon skipping in I-band exons of *TTN*. Whether *TTN* antisense transcript functionality stems from specific nucleotide sequence motifs, RNA secondary structure, or the act of transcription itself remains to be elucidated. However, it is evident from the results of this study that its role in regulation of *TTN* alternative splicing is coupled to RBM20-mediated exon skipping. While it is well established that RBM20 represses exon inclusion in *TTN* and other cardiac genes,^11^ the mechanism by which RBM20 is guided to target pre-mRNA is not well studied. Recently, the splicing factor Srsf1 was shown to be recruited to Triadin (*Trdn*) pre-mRNA via the Triadin antisense transcript (*Trdn-as*) in cardiomyocytes.^17^ Knock out of *Trdn-as* resulted in dysregulated *Trdn* isoform composition, aberrant Ca^2+^-handling and susceptibility to arrythmias. Our results points to a similar mechanism, where *TTN-AS1* facilitates interaction between RBM20 and *TTN* pre-mRNA. While we show that *TTN-AS1* co-localizes with both RBM20 and *TTN* mRNA in cardiomyocyte nuclei, and that knock down of *TTN-AS1* results in decreased interaction between RBM20 and intronic RBM20-binding motifs in *TTN*, elucidating the structural foundation for this mechanism requires further studies. RBM20 interacts with intronic UCUU-motifs in target pre-mRNAs through its RNA recognition motif (RRM).^12^ It is interesting to note that *TTN-AS1-276* has a comparable density of intronic UCUU-motifs (mean of 9.7/kb across all introns) to introns spanning alternatively spliced exons in the RBM20 target genes studied here, *i.e. TTN* (16/kb), *LMO7* (7.3/kb), *CAMK2D* (9.9/kb) and *CACNA1C* (5.1/kb). Based on this observation, we speculate that the interaction with *TTN-AS1* could involve the RRM domain of RBM20.

Knock down of *TTN-AS1-276* in human iPS-CM resulted in a shift towards longer and less stiff sarcomeres with improved diastolic properties. These findings merit further investigation into *TTN* antisense transcripts as therapeutic targets for diseases characterized by increased myocardial stiffness and diastolic dysfunction, such as heart failure with preserved ejection fraction. Genetic models of RBM20 deficiency have been shown to have improved diastolic function^41, 42^ and we believe that *TTN-AS1* could represent a direct target for regulating RBM20 activity through antisense oligonucleotide (ASO) therapies. The 5’ end of *TTN-AS1-276* does not overlap *TTN* exons (nor those of any other gene) and thus represents a feasible target for specific ASO-mediated knock down. Moreover, the fact that *TTN-AS1-276* appears to have its own promoter (**Supplemental Figure 1b**) means that it is amenable to transcriptional modulation by CRISPRi.^43^ Additional studies are required to assess whether interfering with *Ttn* antisense transcription can be harnessed to modulate sarcomere properties and improve cardiac function in *in vivo* models of diastolic dysfunction. One annotated antisense transcript has been identified in the mouse *Ttn* locus (ENSMUST00000156809). As commonly observed among antisense transcripts,^44^, overall sequence conservation with human *TTN* antisense transcripts is low, but it is interesting to note that exon 2 of the mouse transcript has 94% similarity to exon 2 of TTN-AS1-276.

A limitation of the study is the lack of an adult human cardiomyocyte *in vitro* model. iPS-derived cardiomyocytes are known to exhibit a fetal phenotype^45^ and have previously been shown to predominantly express longer N2BA isoforms and a considerably higher N2BA:N2B ratio than adult cardiomyocytes.^36^ However, in our live cell imaging experiments, we observe sarcomere length and contraction dynamics that are comparable to adult cardiomyocytes.

In conclusion, we show that antisense transcripts play an integral role in regulation of *TTN* alternative splicing and sarcomere function in cardiomyocytes and could constitute targets for therapeutic modulation of cardiac stiffness.

## Acknowledgements

This work was supported by grants from the Swedish Heart and Lung Foundation (#20220344, #2023033824 and #2023033924), the Crafoord Foundation and the Royal Physiographic Society. J. Gustav Smith was also supported by grants from the Swedish Research Council (2021-02273), the European Research Council (ERC-STG-2015-679242), Gothenburg University, Skåne University Hospital, governmental funding of clinical research within the Swedish National Health Service, a generous donation from the Knut and Alice Wallenberg foundation to the Wallenberg Center for Molecular Medicine in Lund, and funding from the Swedish Research Council (Linnaeus grant Dnr 349-2006-237, Strategic Research Area Exodiab Dnr 2009-1039) and Swedish Foundation for Strategic Research (Dnr IRC15-0067) to the Lund University Diabetes Center. The authors thank Lund University Bioimaging Centre (LBIC), MultiPark, Center for Translational Genomics and the National Bioinformatics Infrastructure Sweden (NBIS) at SciLifeLab for support.

